# Luxotonic signals in human prefrontal cortex as a possible substrate for effects of light on mood and cognition

**DOI:** 10.1101/2020.09.28.316943

**Authors:** Shai Sabbah, Michael S. Worden, Dimitrios D. Laniado, David M. Berson, Jerome N. Sanes

## Abstract

Animal studies revealed a mood-regulating neural pathway linking intrinsically photosensitive retinal ganglion cells (ipRGCs) and the prefrontal cortex (PFC), involved in the pathophysiology of mood disorders. As humans too have luminance-encoding ipRGCs, we asked whether a similar pathway exist in humans. Here, fMRI was used to identify PFC regions and other areas exhibiting luminance-dependent signals. We report 29 human brain regions where activation either monotonically decreased or increased with luminance. Luxotonic activity was identified across the cerebral cortex, in diverse subcortical structures, and in the cerebellum, regions that have functions related to visual image formation, motor control, cognition, emotion, and reward processing. Light suppressed PFC activation level, the activation monotonically decreasing with increasing luminance. The sustained time course of light-evoked PFC responses, and their susceptibility to prior light exposure, most closely resembled those of ipRGCs. These findings offer a functional link between light exposure and PFC-mediated cognitive and affective phenomena.

## Introduction

Light impacts mood and cognition of humans and other animals. Abnormal lighting induces depression (Bedrosian et al., 2013; Fonken et al., 2012; Gonzalez and Aston-Jones, 2008), while bright light enhances antidepressant therapies and ameliorates both seasonal and non-seasonal depression (Bedrosian and Nelson, 2017; Lam et al., 2016; Penders et al., 2016; Sit et al., 2018). Interestingly, bright light was also found to affect decision-making and working memory, both being key functions of the prefrontal cortex (PFC) (Daneault et al., 2018; Kretschmer et al., 2012; Vandewalle et al., 2007; Xu and Labroo, 2014). These luminance-related effects are thought to depend upon a specialized output channel of the retina dedicated to a stable representation of environmental illumination, constituting a luminance signal (Sabbah et al., 2018; Schmidt and Kofuji, 2009; Wong, 2012). However, the origin of the specific luminance signal acting in the PFC has yet to be identified. One luminance signal revealed in recent years, arises from intrinsically photosensitive retinal ganglion cells (ipRGCs), a rare class of retinal output neurons found both in humans and mice, with autonomous sensitivity to light by virtue of their expression of the photopigment melanopsin (Berson et al., 2002; Ecker et al., 2010). Luminance signals from ipRGCs are conveyed to a myriad of brain regions and affect in mice, among other functions, circadian clock photoentrainment, pupil constriction, neuroendocrine rhythms regulation, learning and mood (Chen et al., 2011; LeGates et al., 2012; LeGates et al., 2014).

Studies using experimental animal models have identified a subset of ipRGCs that transmits light-intensity information to a newly-identified region of the dorsal thalamus, the perihabenular nucleus (PHb); in mice, neurons in the PHb project to the medial PFC (mPFC) (Fernandez et al., 2018), a region known to have a key role in mood regulation (Drevets et al., 2008; Murray et al., 2011). Fascinatingly, manipulation of this retinal-thalamic-frontocortical pathway can induce depression-like behaviors by light in mice (Fernandez et al., 2018). This pathway appears completely separate and radically different from the main retinal-thalamic-visual cortical pathway that targets the visual cortex via the dorsal lateral geniculate nucleus (dLGN). Notably, the PHb also projects to the nucleus accumbens, that modulates hedonic behaviors (An et al., 2020), and the dorsomedial striatum (dmStr), in which activity changes have been associated with goal-directed actions (Graybiel and Grafton, 2015).

These findings imply that in mice, a luminance signal from ipRGCs, is routed to the PFC via a remarkably direct pathway, and that this transduction affects mood, possibly through light-dependent modulation of PFC activity. As humans too have luminance-encoding ipRGCs with highly conserved functional roles (Mure et al., 2019), the critically-relevant question arises – does a similar pathway exist in humans? Studies with humans have indeed revealed links among PFC, light exposure (Penders et al., 2016; Sit et al., 2018) and mood (Baxter et al., 1989; Biver et al., 1994; Brody et al., 2001; Drevets et al., 2002; Drevets et al., 1992; Greicius et al., 2007; Mayberg et al., 1999), and the development of depression and other mood disorders is likely affected by PFC dysfunction (Koenigs and Grafman, 2009; Li et al., 2018; Liotti and Tucker, 1992; Palmer et al., 2014). However, the ability of the human PFC to encode a luminance signal is yet to be addressed. Identifying such a pathway and understanding its function might directly promote development of approaches to treat depression, either by pharmacological manipulation of activation in selected nodes of the pathway, or environmental manipulation with targeted bright light therapy.

Here, as a first step to probing for a retinal-thalamic-frontocortical pathway in humans, we tested the hypothesis that the intensity of light reaching the eyes, stripped of color, form and movement information, would modulate activation in human PFC. We used functional magnetic resonance imaging (fMRI) to identify brain regions having luminance-dependent activation (hereafter ‘luxotonic regions’), and analyzed the data to identify transient and persistent activation modulation. Altogether, 12 brain regions showed steady-state activation according to luminance level. Most were in the PFC or in the classic thalamocortical visual pathway; others were found in the cerebellum, caudate, and the pineal body. PFC and the pineal body exhibited the lowest fMRI signals in bright light, while the other regions exhibited the highest fMRI signal in bright light.

The sustained time course of the light-evoked responses in PFC most closely resembled those of ipRGCs, and prior light exposure influenced the PFC response to light. We also found 17 regions that exhibited transient responses to changes in luminance. These regions were located across the cerebral cortex, diverse subcortical structures, and cerebellum, and they have functions related to visual image formation, motor control, and cognition.

## Results

To determine whether a luminance-dependent functional pathway exists in the human frontal cortex, we used BOLD fMRI (3T Siemens Prisma, 2 mm isotropic voxels) to explore whole-brain activation patterns in 20 healthy, adult participants viewing full-field diffused light stimuli. Every 30 sec, light intensity was stepped to one of four luminance levels. Overall, these spanned nearly four orders of magnitude (10.2, 12.1, 13.1, and 13.8 log photons cm^-2^ s^-1^). Luminance levels were randomized, excluding appearance of a given luminance in consecutive epochs, and under the constraint that all four luminance levels are presented before replication. Each level was presented 15 times (Fig. S1A, B). Participants donned Teflon goggles that diffused the light and removed any spatial contrast, as confirmed by each of the participants. To prevent the participants from falling asleep and closing their eyes, and consequently, limiting the light reaching the retina, the level of alertness was monitored throughout the experiment by asking the participants to discriminate between two randomly-presented tones by pressing one of two buttons (Fig. S1A); all the participants stayed awake with their eyes open throughout the experiment. We used a combination of criteria (monotonic contrast, voxelwise threshold p < 0.001, cluster false discovery rate = 0.05, cluster size > 27 voxels) to reveal brain regions that exhibit activation that varied monotonically with luminance and show either a sustained or transient component (Fig. S1C, D, see *Methods* for details).

### Luminance modulates steady-state activation of diverse cortical, subcortical, and cerebellar regions

Overall, 12 brain regions met the criteria for luminance-dependent monotonic activation, with a significant sustained component (see *Methods* for details on the regressor used for the analysis) (Fig. 1A). Luminance-dependent activation in all these regions persisted throughout the 30-sec duration of the stimulus. In several of the regions, activation also changed abruptly upon light onset (Fig. 1B, Fig. S2). Steady-state BOLD signals in those 12 brain areas with a sustained component either monotonically increased or decreased with luminance level. Six of these regions were located in the PFC, specifically, in the anterior prefrontal, orbitofrontal, and anterior cingulate cortices (regions S1-S6), all showing a monotonic *decreasing* response to increasing luminance. By contrast, monotonically *increasing* responses to increasing luminance were observed in the image-forming visual system: light intensity strongly increased the BOLD signal in the occipital lobe (S7), especially in the primary visual cortex (V1), but also in a number of extrastriate regions beyond V1. These results are consistent with previous studies (Boyaci et al., 2007; Haynes et al., 2004; Vinke and Ling, 2020), but see (Cornelissen et al., 2006). The LGN (S8, S9), the thalamic relay nucleus of the primary visual pathway, is almost certainly the source of this luxotonic signal in V1, since it too exhibited bilateral, strong BOLD activation responses to bright light, that monotonically increased with luminance level. The only brain region beyond the primary visual pathway that shared the visual system’s BOLD activation to light was a small region in the uvula at the inferior vermis of the cerebellum (S12). Two additional small regions exhibited suppressed BOLD responses that monotonically decreased with luminance, like those in the PFC. These were the right caudate nucleus (S10) and the pineal body (S11), in which activity is known to be depressed in response to light through the influence of ipRGCs (Hull et al., 2018). Sensitivity, calculated as the difference in steady-state response between the lowest and highest stimulus luminance divided by the change in luminance, was significantly larger in cortical (0.12±0.04 BOLD response; mean±s.d.) compared to subcortical regions (0.04±0.02 BOLD response; permutation t-test, p = 0.006) (Fig. S2K).

**Figure 1.**
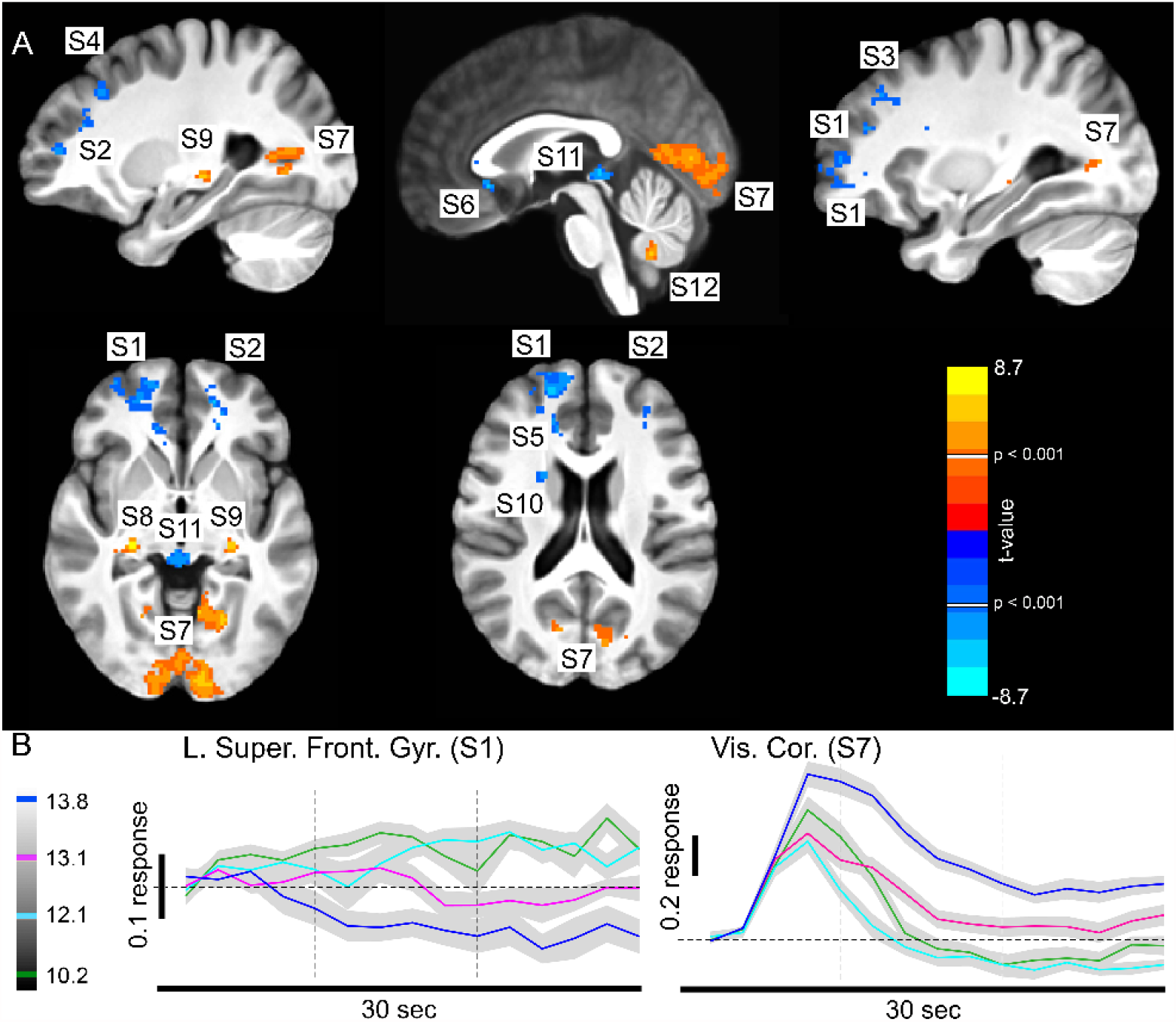
Cortical, subcortical, and cerebellar luxotonic regions showing a sustained component. **(A)** The 12 identified luxotonic regions are color-coded based on the contrast for monotonic increases or decreases as a function of luminance level. Color scale corresponds to *t*-values: warm colors (positive t) represent light-evoked responses increases with luminance level; cool colors (negative t) depict light-evoked responses decreases with luminance level. Color bar, *t*-values range ±8.7, and images are thresholded at p < 0.001, t(two-tailed) = 3.621 (white horizontal lines). Anatomical underlay represents the mean of all 20 subjects following spatial alignment. **(B)** BOLD responses for S1 and S7 over the 30-sec light stimulus, for four luminance levels. Data for all luxotonic regions are shown in Fig. S2.

### Functional connectivity between luxotonic sustained regions

We next assessed to what extent the activated regions formed independent or dependent networks, by assessing functional connectivity between each of the 12 regions that exhibited sustained luminance-dependent activation (Fig. 2A). BOLD time-series in luxotonic regions in the frontal cortex generally exhibited significant positive correlations with one another (Fig. 2A, hot colors), except for the left middle frontal gyrus which is located posteriorly to all the other analyzed prefrontal regions. Similarly, we found positive correlations in the BOLD time-series among the regions along the thalamo-visuocortical pathway: V1 (with extra striate areas) and LGN as well as a region in the vermis of the cerebellum. These BOLD time-series were either not correlated or negatively correlated with prefrontal cortical regions (Fig. 2A, cold colors). Thus, it seems that sustained luminance-dependent responses engaged two main separate functional networks, one encompassing the thalamo-visuocortical pathway and another including a set of prefrontal cortical regions. The latter, were previously implicated in mood regulation and high-level cognition and value-related judgments (Daneault et al., 2018; Kretschmer et al., 2012; Vandewalle et al., 2007; Xu and Labroo, 2014).

**Figure 2.**
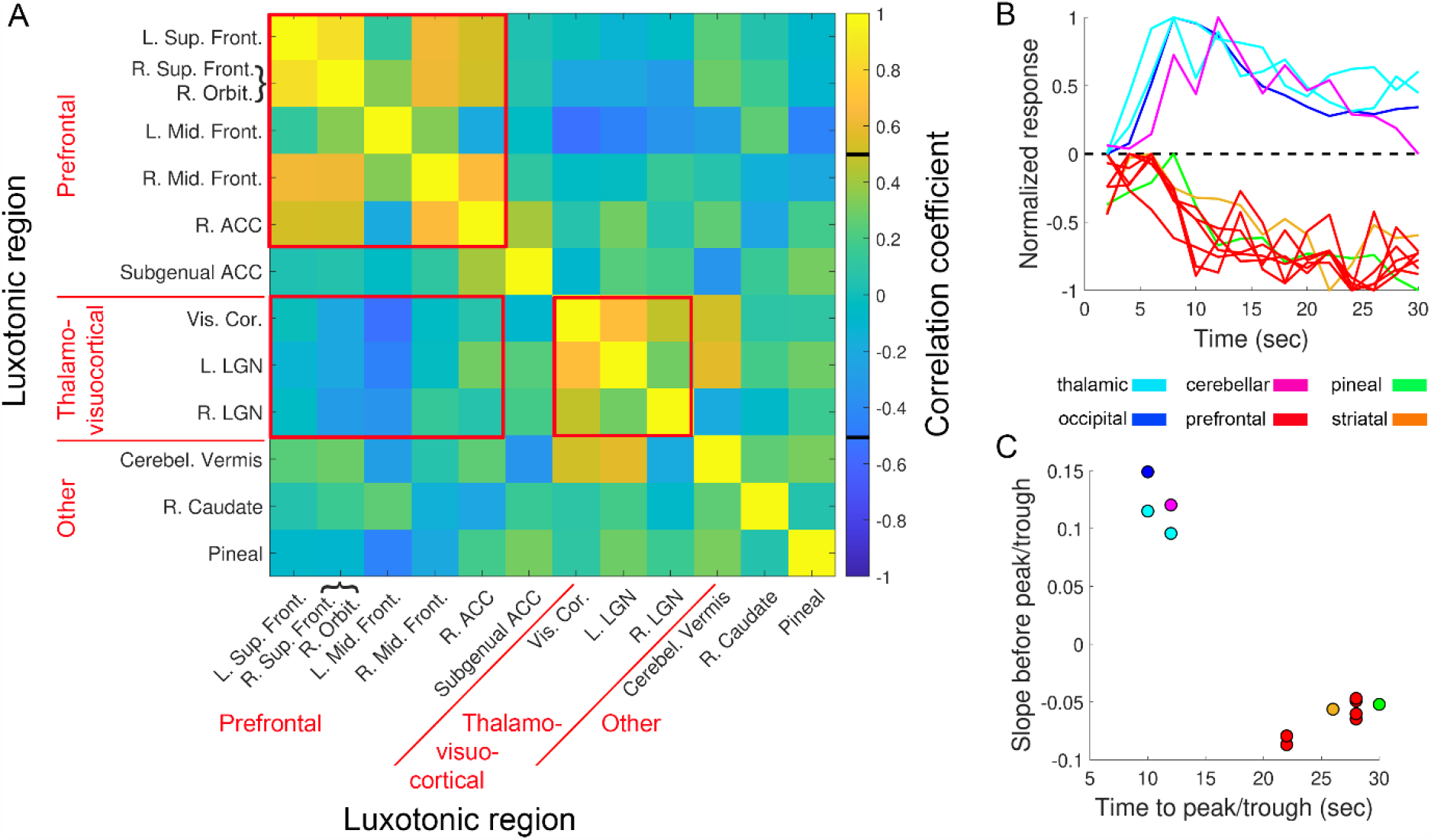
Light-evoked responses with a significant sustained component are either rapid and increasing with luminance or slow and decreasing with luminance. **(A)** Pair-wise Pearson correlation between the 12 identified luxotonic regions. Regions are grouped as ‘Prefrontal’, ‘Thalamo-visuocortical’, and ‘Other’. Correlation coefficient of ±0.51 corresponds to a significance level of 0.05 (indicated on the color scale bar). **(B)** Light-evoked responses, normalized in all 12 identified luxotonic regions to the highest-luminance level, averaged across 20 participants, fell into two functional groups: rapid and slow. Responses are color-coded based on the general brain region, as depicted in the legend at the bottom. **(C)** Distribution of the 12 identified luxotonic regions as a function of the slope before peak/trough and the time to peak/trough. Responses are color-coded based on the general brain region.

### Activation in luxotonic sustained regions diverges in temporal profile and polarity

Our findings represent the first demonstration that frontal cortical areas, and the cerebellar inferior vermis, caudate nucleus, and pineal body, exhibit activation modulation in proportion to luminance level. To start studying the timing of the sustained light-evoked BOLD responses, we compared them by normalizing the amplitude of all observed time courses to range between 0 and 1 (or -1) (Fig. 2B). Luxotonic regions in the visual cortex, LGN, and cerebellum, in which light enhanced the BOLD signal, showed a relatively rapid (latency 10 sec) increase in activation, which then gradually declined in magnitude (‘rapid set’, n = 4) (Fig. 2B, blue and cyan lines). In contrast, activity in all the prefrontal luxotonic regions, caudate nucleus and pineal body, activity dropped below baseline reaching its minimum toward the end of the 30-sec stimulus presentation (‘slow set’, n = 8) (Fig. 2B, red, magenta, green and orange lines). This dichotomy in the temporal profile and polarity of the responses is clearly evident when plotting the time to peak/trough as a function of the slope before peak/trough (Fig. 2C).

### Effect of light history on prefrontal luxotonic activation

Evidence demonstrates that prior light exposure can modulate subsequent brain activity and the emotional and cognitive processes mediated by the specific brain region (Alkozei et al., 2016b; Shan et al., 2015; Vandewalle et al., 2006). Thus, to remove the effect of prior light exposure on the ability of the 12 identified luxotonic regions to encode luminance, we focused on the three transitions in which luminance level (L) was stepped up from the lowest intensity (L1 → L2, L1 → L3, and L1 → L4). This revealed that the prefrontal activation in response to the second stimulus in each luminance pair (L2, L3, and L4) developed slowly throughout the 30-sec duration of the stimulus (Fig. 3, left panels). Steady-state prefrontal activation, measured as the average response over the last 10 sec of the 30-sec stimulus, monotonically decreased with increasing luminance level in all six identified prefrontal luxotonic regions (Fig. 3, panels on second column from the left), namely, the left and right superior frontal gyri (S1,S2), right orbital gyrus (S1), left and right middle frontal gyri (S3,S4), right anterior cingulate cortex (S5), and subgenual anterior cingulate cortex (S6). In addition to the six prefrontal regions, also the pineal body (S11) and right striatal caudate (S10) encoded steady-state luminance level. Activation in these regions was similarly monotonically suppressed with increasing luminance (Fig. S3A,B). In contrast, as Fig. 4A shows, the effect of light exposure on activation along the thalamo-visuocortical pathway was different. In the visual cortex (S7), activation increased sharply several seconds after the light onset, and then gradually rolled off, never returning to baseline during the 30-sec light stimulus (Fig. 4A, left column). Similar trends were observed in the left (S8) and right (S9) LGN (Fig. 4B,C, left column). While steady-state activation increased with luminance in all three identified regions along the thalamo-visuocortical pathway (Fig. 4A-C, second column from left), the magnitude of responses in the visual cortex was twice as large as that observed in the LGN. Activation in another luxotonic region, the cerebellar vermis (S12), also appeared to slightly increase with increasing luminance (Fig. S3C).

**Figure 3.**
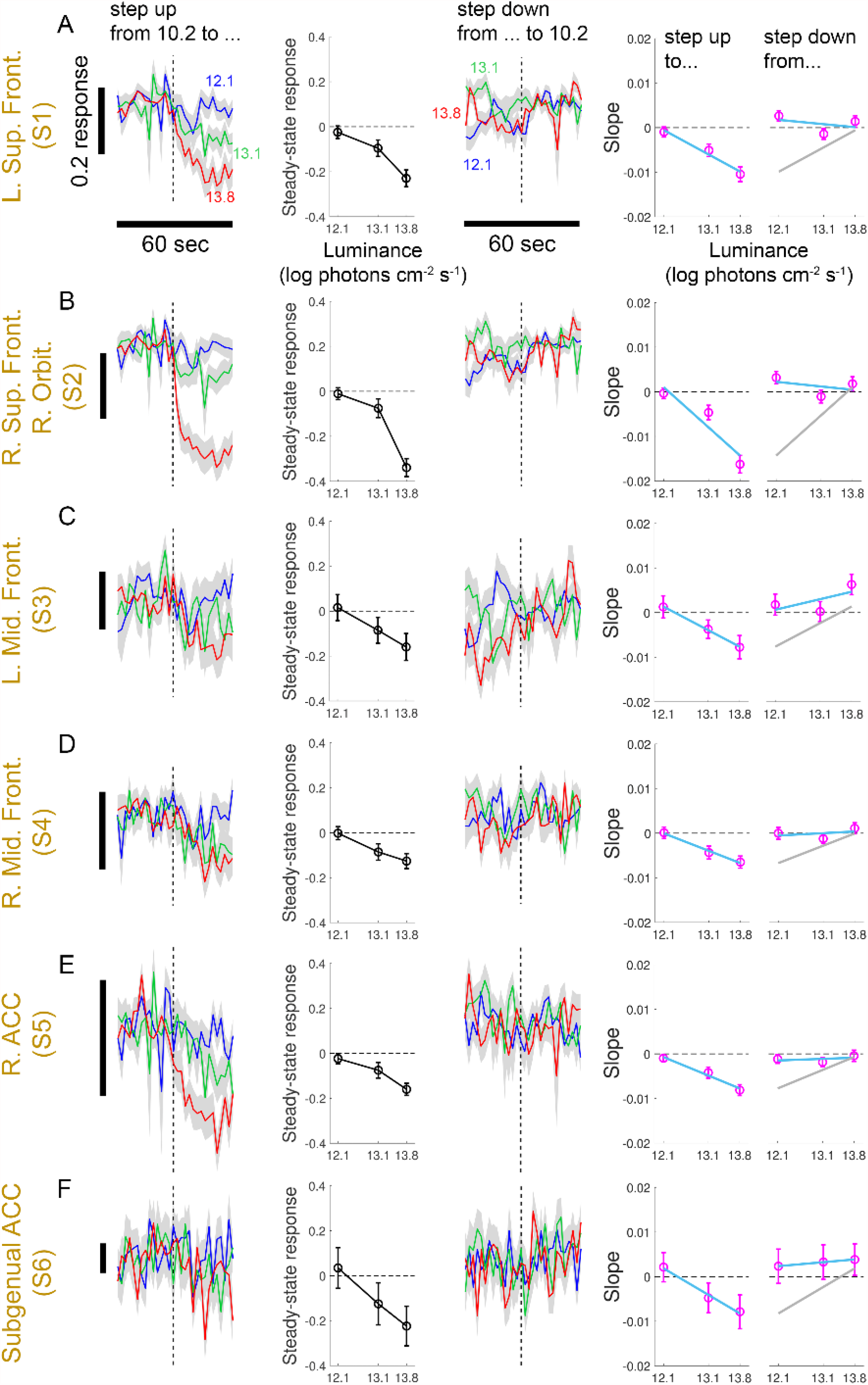
Light suppresses steady-state prefrontal activity. **(A-F) left.** BOLD Light-evoked responses to step up transitions – transitions in which luminance was stepped up from the same, lowest, luminance level. Responses around individual transitions are depicted in different colors. Shaded gray areas represent the standard error from the mean. Dashed vertical line marks the transition time. For example, the red curve in panel A represents stepping up luminance from the lowest level (L1, 10.2 log photons cm^-2^ s^-1^) to the highest level (L4, 13.8 log photons cm^-2^ s^-1^). (**A-F**) **second column from left**. Steady-state prefrontal activity monotonically decreased with increasing luminance level in all six identified prefrontal luxotonic regions. These are the left and right superior frontal gyri and right orbital gyrus (A,B), left and right middle frontal gyri (C,D), right anterior cingulate cortex (E), and subgenual anterior cingulate cortex (F). Vertical bar representing 0.2 BOLD response in *A* applies to the amplitude of responses in *A*-*F*. (**A-F**) **third column from left**. BOLD Light-evoked responses to step-down transitions (transitions in which luminance was stepped down from the same, highest, luminance level). (**A-F**) **right**. Slopes fitted to the three step-up and three step-down transitions. In the prefrontal cortex in which activity decreased with luminance level, transitions to higher luminance levels resulted in positive slopes, whereas transitions to lower luminance levels resulted in negative slopes. With no effect of light history, the trend observed when moving across slope values for step-up transitions (cyan line) is expected to be the opposite of that of step-down transitions (gray line). The actual observed trend for step-down transitions is depicted (cyan line). The angular difference between the observed and expected trends for step-down transitions serves as a proxy for the strength of the effect of prior light history. Sample size, L1 → L2: n = 81, L1 → L3: n = 56, L1 → L4: n = 79, L2 → L1: n = 78, L3 → L1: n = 84, L4 → L1: n = 68.

**Figure 4.**
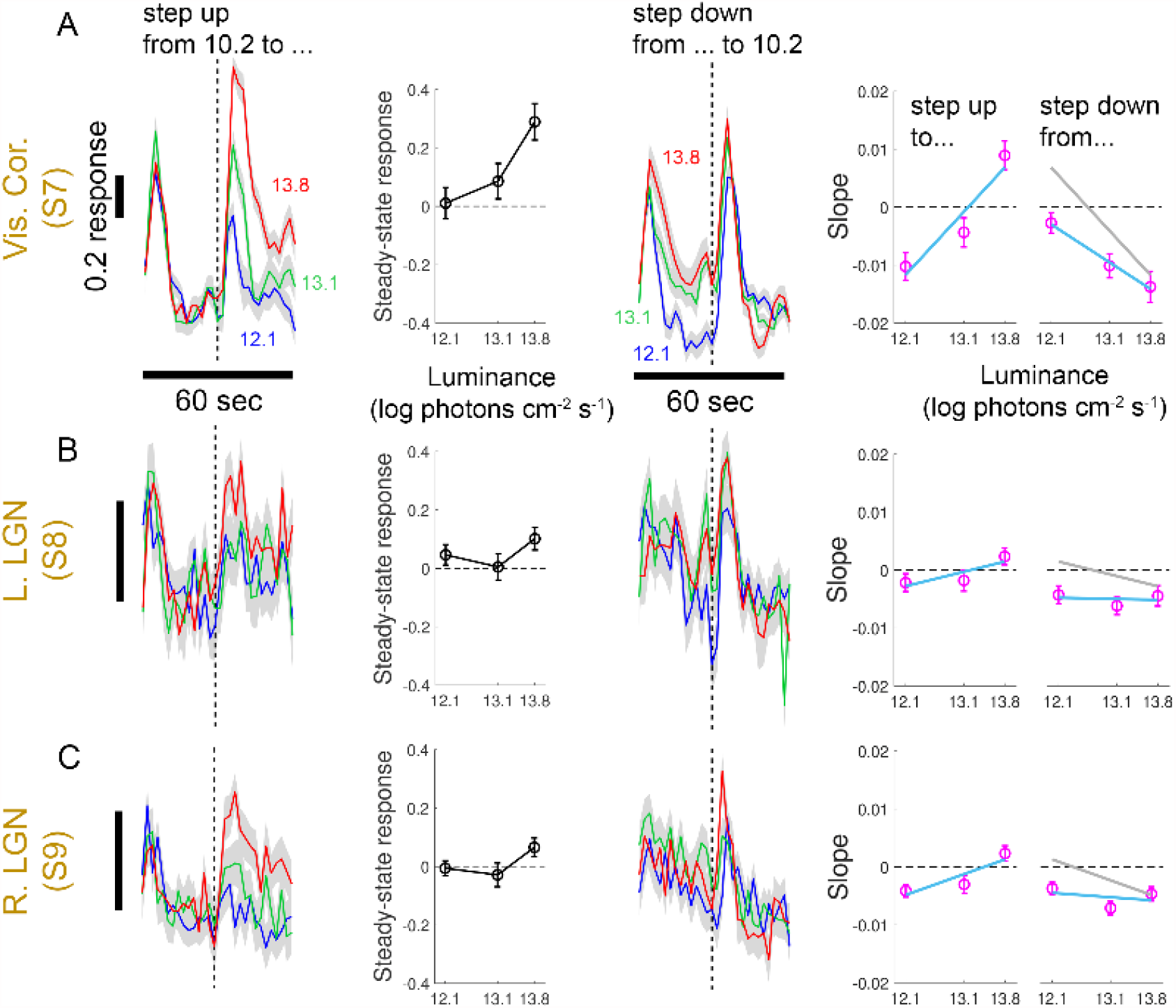
Light enhances activity along the thalamo-visuocortical pathway. Conventions as in Fig. 4. (A-C, two leftmost columns). Steady-state activity monotonically increased with increasing luminance level in all three luxotonic regions along the thalamo-visuocortical pathway. Vertical bar, representing 0.2 BOLD response in *A*, applies to all panels. In the visual cortex (*A*) and the LGN (*B,C*), in which activity increased with luminance level, transitions to higher luminance levels resulted in zero or negative slopes, whereas transitions to lower luminance levels resulted in zero or positive slopes. Sample size, L1 → L2: n = 81, L1 → L3: n = 56, L1 → L4: n = 79, L2 → L1: n = 78, L3 → L1: n = 84, L4 → L1: n = 68.

Next, to quantify the effect of prior light exposure on BOLD activation, we plotted the three transitions in which luminance level was stepped down from the highest luminance level, that is, L4 → L1, L3 → L1, and L2 → L1 (Fig. 3, third column from left). We then fit a straight line to the activation around a given transition and repeated this for the three step-up (Fig. 3, left panels) and the three step-down (Fig. 3, third column from left) transitions. This yielded three slope values for step-up transitions and three slope values for step-down transitions (Fig. 3 right). If brain activation is not affected by prior light exposure, the trend observed when moving across slope values should be similar for the step-up and step-down transitions. Deviation from this expectation serves as a proxy for the strength of the effect of prior light exposure. While activation in all 12 identified luxotonic regions was affected by prior light exposure, the effect was largest in the right and left superior frontal gyri and subgenual anterior cingulate gyrus (Fig. 5; Fig. 3, right; Fig. 4, right; Fig. S3, right). These results demonstrate a strong effect of light history on prefrontal activation. ipRGCs, more so than conventional retinal ganglion cells, can exhibit persistent responses following the light offset (Pang et al., 2003). Therefore, this analysis raises the possibility that the observed PFC activation, at least in the right and left superior frontal gyri and subgenual anterior cingulate gyrus, is consistent with ipRGCs effects.

**Figure 5.**
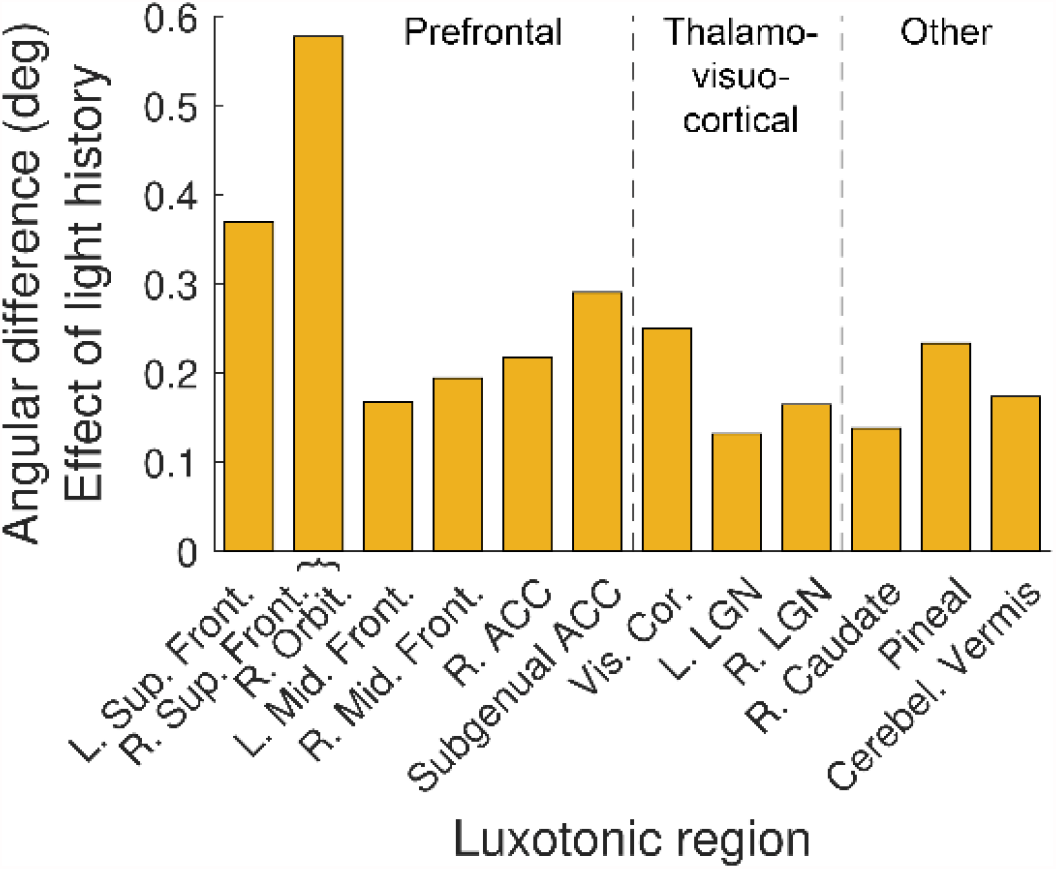
The effect of prior light exposure on brain activity in the prefrontal cortex. The magnitude of the effect of light history estimated as the angular difference between the observed and expected trends for step-down transitions. While prior light exposure affected all luxotonic regions, the effect on the left and right superior frontal gyri and right orbital gyrus was the largest.

### Luminance induced transient activation in the cerebral cortex, subcortex, and cerebellum

In addition to the 12 luxotonic regions with a significant sustained component that we identified above, we also identified 17 regions that exhibited activation with a significant transient component (see *Methods* for details on the regressors used for this analysis). Also in these regions, activation was luminance-dependent; transient activation increased (orange) or decreased (blue) monotonically with luminance (Fig. 6). Luminance-dependent transient activation was detected bilaterally in the superior medial frontal gyri (T1, T2), precentral gyri (T6, T7), visual cortex T13), and paracentral lobule (T11). Right-hemisphere-only transient activation was detected in the supplementary motor area (T5), frontal operculum (T8, T9), parietal operculum (T10), and insula (T12). And, left-hemisphere-only transient activation was detected in the middle frontal (T3), anterior cingulate (T4), middle occipital (T14), and inferior occipital (T15) gyri, and cerebellum (T16, T17).

**Figure 6.**
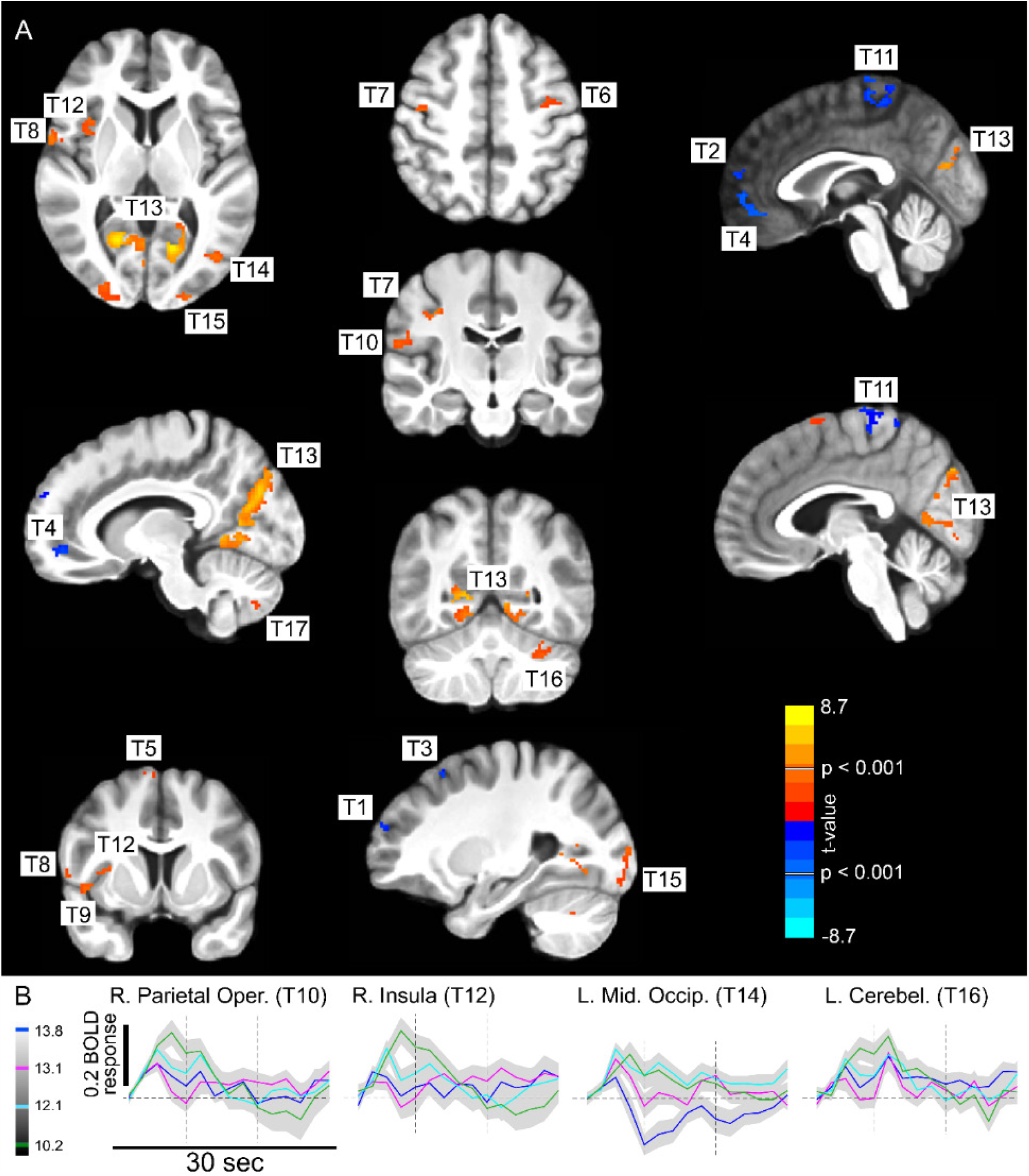
Cortical, subcortical, and cerebellar luxotonic regions that show a significant transient component. (**A**) The 17 identified luxotonic regions are color-coded based on the contrast for monotonic increases or decreases as a function of luminance level. Color scale corresponds to t-value: warm colors (positive t) – light-evoked responses increases with luminance level; cool colors (negative t) – light-evoked responses decreases with luminance level. Color bar, t-values ranges ±8.7, and images are thresholded at p < 0.001, t(two-tailed) = 3.621 (white horizontal lines). Anatomical underlay represents the mean of all 20 subjects following spatial alignment. Regions: T1, L. Super. Medial Front.; T2, R. Super. Medial Front.; T3, L. Mid. Front.; T4, L. ACC; T5, R. Suppl. Motor Area; T6, L. Precentral; T7, R. Precentral; T8, R. Front. Oper.; T9, R. Front. Oper.; T10, R. Parietal Oper.; T11, Paracentral Lob.; T12, R. Insula; T13, Vis. Cor.; T14, L. Mid. Occip.; T15, L. Inf. Occip.; T16, L. Cerebel.; T17, L. Cerebel. (**B**) BOLD responses for T10, T12, T14, and T16 over the 30-sec light stimulus, for four luminance levels. Data for all the luxotonic regions are shown in Fig. S4.

We next assessed whether the 17 luxotonic regions exhibiting a transient component, formed independent or dependent networks by assessing their functional connectivity (Fig. 7). BOLD time-series in the frontal cortex showed two distinct activation sets, one in the PFC (Set ***a***) and another in more posterior regions including motor areas (Set ***b***). Time-series in Set ***b*** correlated with those in the parietal operculum. Time-series in the paracentral lobule (Set ***c***) positively correlated with those in the prefrontal clusters but negatively correlated with all other regions. Time-series in the left inferior occipital gyrus did not correlate with any of the other regions (Set ***d***). Time series in the visual cortex and left middle occipital gyrus correlated with one another.

**Figure 7.**
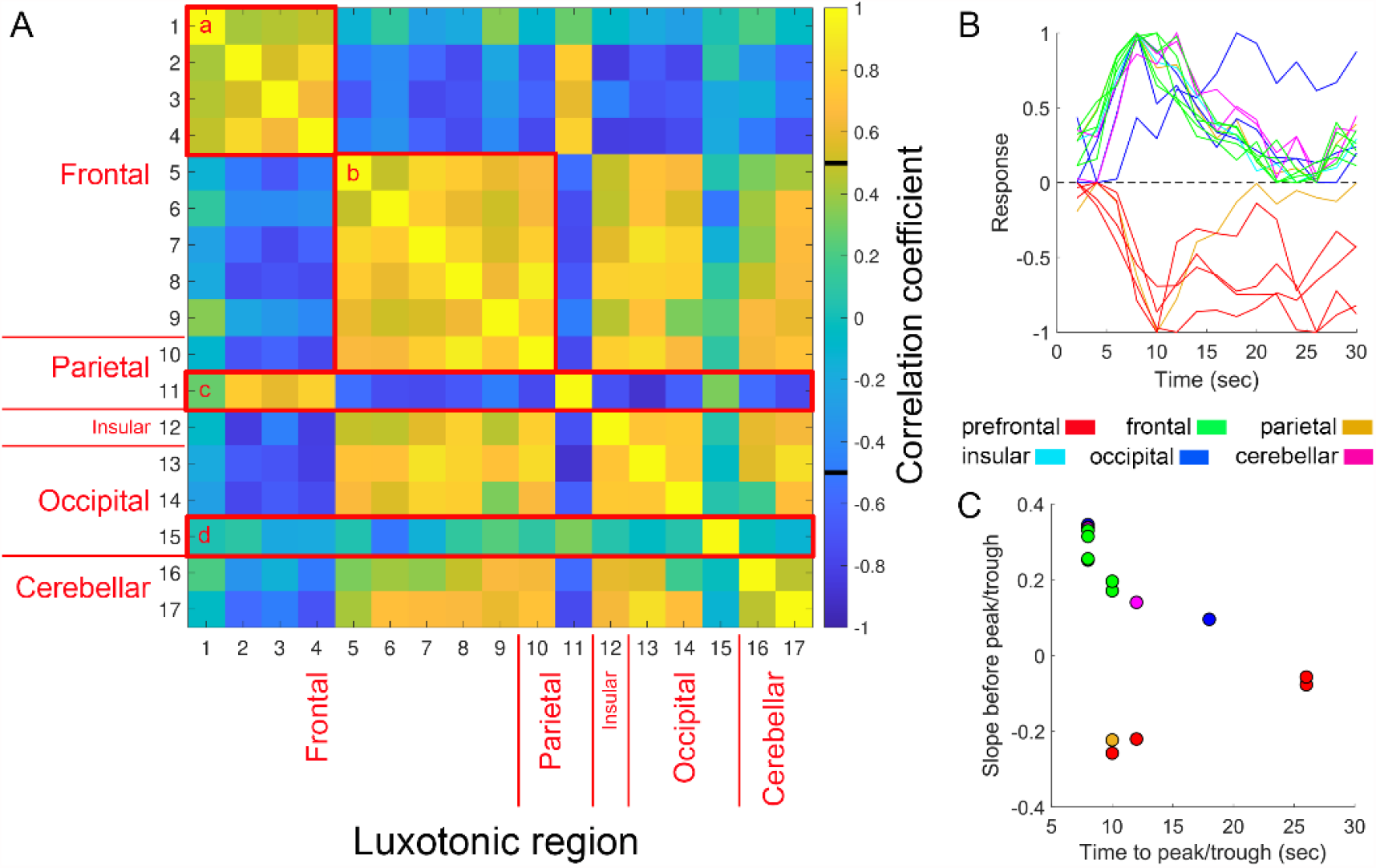
Pair-wise functional connectivity between identified luxotonic regions with a transient component (A) Pair-wise Pearson correlation between the 17 identified transient luxotonic regions. Regions are grouped as frontal, parietal, insular, occipital, and cerebellar. Identity of luxotonic brain regions is given in Fig. 6. A correlation coefficient of ±0.51 corresponds to a significance level of 0.05 (indicated on the color scale bar). **(B)** Normalized light-evoked responses to the highest-luminance level in all 17 identified luxotonic regions, averaged across 20 participants, fell into two main functional groups: increasing and decreasing, but otherwise varied considerably in the steady-state phase. Responses are color-coded based on the general brain region, as depicted in the legend at the bottom. **(C)** Distribution of the 17 identified luxotonic regions as a function of the slope before peak/trough and the time to peak/trough. Responses are color-coded based on the general brain region.

Next, just as we did for luxotonic regions in which activation exhibited a sustained component (Figs. 3 and 4, Fig. S3), we assessed the luminance encoding capability of the 17 identified luxotonic regions that exhibited a transient component while controlling for prior light exposure – we examined the three transitions in which the luminance level was stepped up from the lowest intensity (Fig. S4). Luminance-dependent activation level typically changed abruptly upon light onset, and in several regions, persisted throughout the 30-sec duration of the stimulus. Activation in the superior medial and middle frontal gyri as well as in the anterior cingulate cortex (regions T1-T4) decreased sharply following light onset, remained relatively low throughout the 30-sec duration of the stimulus, and typically decreased monotonically with increasing luminance.

Activation in the 13 remaining identified regions increased upon light onset and then gradually rolled off. In some regions (*e*.*g*., visual cortex, T13) activation did not return to baseline during the 30-sec light stimulus.

### Luxotonic regions exhibiting sustained and transient activation co-occur along a streak in the occipital lobe

Finally, we examined the overlap between the group of luxotonic regions with a sustained component and the group with a transient component. Overlap was not detected except for one nearly contiguous strip running along the parieto-occipital sulcus and the dorsal cuneus (Fig. 8), indicating that sustained and transient responsive regions were almost completely segregated. Sustained activation comprised 4,766 voxels and transient activation comprised 3,403 voxels, for a total of 7,484 voxels. Of these, 685 voxels had an overlap for transient and sustained activation, corresponding to ∼9% of the combined activation.

**Figure 8.**
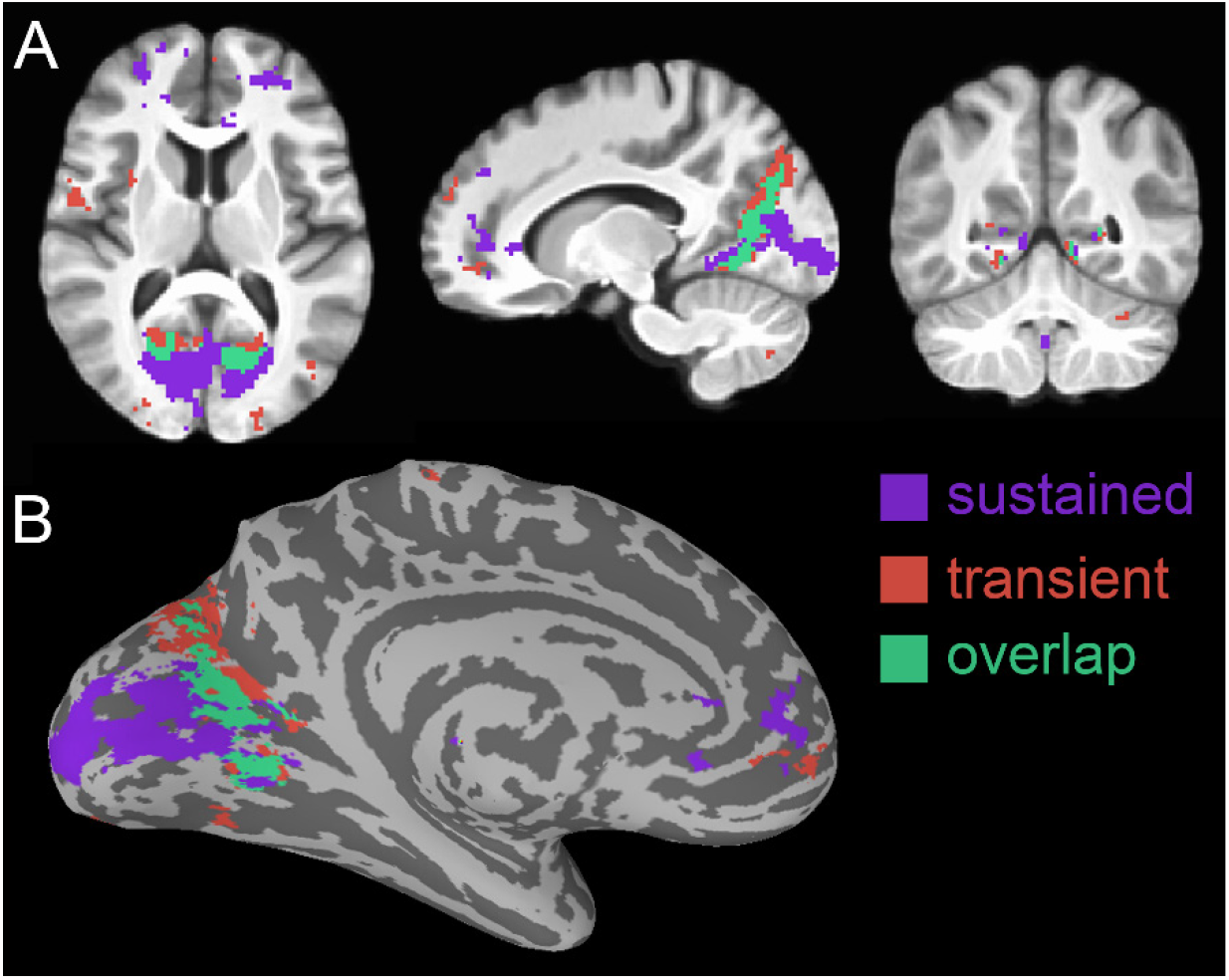
Sustained and transient response regions along the parieto-occipital sulcus. (**A**) Horizontal, mid sagittal and coronal sections showing the overlap between luxotonic regions in which activity showed both significant sustained and transient components. Sustained and transient regions are marked in purple and in red, respectively, and overlapping areas are marked in green. (**B**) Same data presented on an inflated brain model.

## Discussion

While luxotonic activation in the human visual cortex has been previously documented (Boyaci et al., 2007; Haynes et al., 2004; Vinke and Ling, 2020), to the best of our knowledge, this is the first report of luxotonic activation in the human prefrontal cortex, precentral gyri, paracentral lobule, frontal and parietal operculum, insula, supplementary motor area, LGN, caudate nucleus, pineal, and cerebellum. Therefore, our results demonstrate the effects of light on diverse human brain regions that collectively contribute to motor control, cognition, emotion, and reward processing. We identified 29 human brain regions in which activation either monotonically decreased or increased with luminance. Activation in 12 regions exhibited a significant sustained component, whereby 17 other regions exhibited a significant transient activation pattern. All the regions in which activation decreased monotonically with increasing luminance showed responses that persisted until the termination of the light stimulus; in several of them the effect of light persisted even longer.

### Acute effect of light on steady-state PFC activation

Prior observations have suggested light-evoked modulation of several prefrontal regions. For example, blue light exposure (as opposed to green or orange) concurrently with an auditory working memory task, increases activation in the left middle frontal gyrus (Daneault et al., 2018; Vandewalle et al., 2007) and decreases anterior cingulate cortex (ACC) activation (Daneault et al., 2018). Furthermore, blue light (as opposed to amber), decreases ACC activation following an emotional anticipation task (Alkozei et al., 2016a). However, our observations uniquely demonstrate wide-ranging effects of light on PFC physiology, which varied monotonically with luminance level. We identified six luxotonic regions having a sustained-activation pattern in the PFC: right and left superior frontal gyri, right and left middle frontal gyri, right anterior cingulate cortex, and subgenual anterior cingulate cortex. These appear to be involved in cognitive and emotional processes. For example, the left superior frontal gyrus is involved in self-awareness (Goldberg et al., 2006) and spatial working memory (Courtney et al., 1998). In the right hemisphere, this region contributes to the control of impulsive response or action inhibition (Hu et al., 2016). The middle frontal gyrus has a role in regulating working memory (Nee et al., 2013) and reorienting attention (Japee et al., 2015), and has been designated as a component of the Ventral Attention Network (Corbetta et al., 2000) – a bottom-up system that responds to unexpected stimuli in an involuntary manner (Corbetta et al., 2008). The ACC contributes to social cognition, attention allocation, motivation, decision making, learning, cost-benefit calculation, impulse control, and emotion (Ridderinkhof et al., 2004; Shackman et al., 2011).

And, finally, the subgenual ACC is associated with the regulation of emotions and the mechanism of action of antidepressants (Drevets and Savitz, 2008). Our results demonstrate that activation in all six identified prefrontal luxotonic regions decreased with luminance, and offer a possible functional link between frontal light processing and a number of cognitive and affective phenomena previously reported to be affected by light intensity.

### Chronic effect of light on prefrontal activation

In addition to the acute effect of light on neural activation, light can also affect neural activation chronically. We found a pronounced effect of light exposure on subsequent activation in the right and left superior frontal gyri and the right orbital gyrus. Activation in those regions, which comprise portions of the medial PFC (mPFC), was suppressed during light exposure as well as following it (Fig. 5C). Such chronic suppression of neural activation by light has not been previously reported in these regions, but chronic enhancement of neural activation by light have been reported in other brain regions. For example, a single, 30-min exposure to blue as opposed to amber light (expected to activate the blue-sensitive melanopsin photopigment of ipRGCs) has been shown to improve working memory task performance and increase neural activation in the dorsolateral (dlPFC) and ventrolateral (vlPFC) PFC over the subsequent 40-min period (Alkozei et al., 2016b). Another study, using a shorter light exposure (21 min.), reported increased activation over the subsequent 8-min. period in the hippocampus, right anterior cingulate cortex, left precuneus, and right intraparietal sulcus (Vandewalle et al., 2006). Our demonstration of a chronic effect of light on PFC activation complements previous reports and may constitute the substrate for the previously reported effects of light on decision making, social behavior, working memory, and mood (Alkozei et al., 2016b; Bedrosian et al., 2013; Knez, 2001; Shan et al., 2015; Wessolowski et al., 2014).

### Origin of prefrontal luxotonic responses

The slowly-evolving and persistent responses we observed in the human PFC are reminiscent of ipRGCs responses (Fig. 2A, 3). ipRGCs responses in humans (Mure et al., 2019), as well as in the mouse (Estevez et al., 2012; Sabbah et al., 2018; Schmidt and Kofuji, 2009; Wong, 2012), persist throughout the duration of the stimulus and even after light offset, with the post-stimulus response duration increasing with luminance. Thus, the similarity in light-evoked activity in ipRGCs and the human PFC may suggest a contribution of ipRGCs’ light responses to PFC activation. While ipRGCs receive synaptic input from rod/cone photoreceptors, their melanopsin pigment is especially sensitive to blue light wavelengths (Berson et al., 2002) that have been shown to shape PFC-mediated functions in humans. For instance, blue-enriched white light occurring in the evening (as opposed to white light), either in the environment or emitted by a computer screen, improves cognitive performance, alertness, and concentration (Cajochen et al., 2011; Viola et al., 2008). Improvement in cognitive performance under blue-enriched lighting has been reported also during the day (Lehrl et al., 2007). At least some of these cognitive functions have been shown to depend on ipRGCs, tying specific frontal functions to a unique ipRGCs sensitivity. For example, in blind individuals who lack rod/cone photoreceptors but retain ipRGCs, blue light exposure during an auditory task has been shown to stimulate activation in the medial PFC and ventrolateral PFC (Vandewalle et al., 2013), supporting the notion that the light-evoked modulation we observed in the PFC had contribution of ipRGCs. Together, these reports suggest that alteration in cognition and frontal brain activation is partly driven by blue-light-sensitivity, and specifically by ipRGCs as the transducers of this sensitivity.

By contrast, the rapidly-developing and -terminating responses we observed in the human V1 and LGN, resemble responses of conventional RGCs (non-ipRGCs) – they gradually roll off and terminate at light offset. Conventional RGCs do not express melanopsin, they respond to alterations in light intensity only briefly, and their maintained firing rates are not correlated with photon flux (Baden et al., 2016; Sabbah et al., 2017). Conventional RGCs encode temporal and spatial contrast, and support the detection and recognition of objects, color, depth and motion (Sanes and Masland, 2015; Wei et al., 2011), especially via their signals routed through the thalamus to the visual cortex. Interestingly, the luxotonic activity we observed in V1 and LGN did not return to baseline by stimulus termination, suggesting, albeit without proving, a modest ipRGCs’ contribution to this activation pattern. Indeed, melanopsin, the blue-sensitive pigment of ipRGCs, have been shown to contribute to light-evoked response in the human V1 (Spitschan et al., 2017); and, while the majority of input to the mouse dLGN, which routes information to visual cortices, arrives from conventional RGCs, at least some ipRGC types do project to the dLGN (Hannibal et al., 2014). Therefore, the light-evoked activation we observed in the visual cortex and LGN likely represents the integration of signals from both ipRGCs and conventional RGCs.

Our results suggest no clear correspondence between the luxotonic regions in humans and those in mice, though differences in homologous pathways could readily explain discrepancies. In mice, the thalamic PHb projects to the infralimbic and prelimbic cortices (Fernandez et al., 2018), raising the possibility that luxotonic neural input is transmitted from the PHb to those limbic cortices. However, we did not find luminance-dependent activation in the infralimbic and prelimbic cortices of humans. This may have resulted from our conservative criterion for minimum cluster size (27 voxels), the limited size of the human infralimbic and prelimbic cortices, or from differences in homologous pathways. Nonetheless, while it is currently unknown whether the human PFC is part of a pathway homologous to the retinal-thalamic-frontocortical pathway discovered in mice, the persistence and luxotonic nature of light-evoked responses in the human PFC strongly suggest that it receives input from ipRGCs, similarly to the pathway in the mouse (Fernandez et al., 2018). Moreover, mouse PHb neurons that receive ipRGCs input, as well as PHb neurons that project to the NAc, have been shown to be mainly GABAergic (An et al., 2020). If at least a subset of mPFC-projecting PHb neurons is also GABAergic, this would raise the possibility that the mouse PHb transmits an inhibitory drive to the PFC. If discovered in humans, such PHb-to-mPFC inhibitory input may explain our observation that light exposure suppresses prefrontal activation.

### Human PFC and mood regulation

The PFC is commonly divided, based on connectivity and functional specialization, into the dorsolateral (dlPFC) and ventromedial (vmPFC) (Koenigs and Grafman, 2009; Kringelbach and Rolls, 2004). Resting state brain activation (Biver et al., 1994; Drevets et al., 1992) and glucose metabolic rates (Biver et al., 1994; Drevets et al., 2002) in vmPFC, including the subgenual ACC and orbitofrontal gyrus, are higher in depressed people, whereas the metabolic rates in dlPFC of depressed people appear low (Baxter et al., 1989; Biver et al., 1994). Recovery from depression and antidepressants affect the vmPFC and dlPFC in an opposite manner. Specifically, antidepressants and the accompanying recovery from depression is associated with decreased vmPFC activation but increased dlPFC activation (Brody et al., 2001; Mayberg et al., 1999). Here, we found light-evoked suppression of PFC, including in the subgenual ACC and orbitofrontal gyrus. Therefore, similar to antidepressants, light appears to reduce activation in these regions.

### Effect of light on the pineal, cerebellum, and caudate nucleus

We identified luxotonic regions with persistent activity in the pineal, cerebellum, and caudate nucleus. Activation in the caudate nucleus was suppressed in a luminance-dependent manner. While the functional role of this luminance-dependent modulation is unknown, it raises the possibility that light might affect the reward system, motor control, procedural and associative learning, and inhibitory control of action (Grahn et al., 2008). Indeed, long-term blue light exposure has been shown to enhance risk-related activation in the ventral striatum and head of caudate nucleus (Macoveanu et al., 2016).

We identified a luxotonic region in the uvula that constitutes a large portion of the cerebellar inferior vermis. Activation in this region was enhanced with light exposure. The uvula is being activated during optokinetic eye movements, and this activation depends on the speed and direction of eye movements (Ruehl et al., 2017). Interestingly, oculomotor performance has been shown to depend on luminance, resulting in eye movements of greater accuracy in the light (Goffart et al., 2006), suggesting a possible functional role for the uvula’s luminance-dependent activation in oculomotor responses.

The melatonin-producing pineal gland modulates sleep and circadian and seasonal rhythms (Macchi and Bruce, 2004). It receives indirect photic input from ipRGCs (via the paraventricular hypothalamic nucleus, intermediolateral nucleus of the spinal cord, and superior cervical ganglion) (Borjigin et al., 2012). Our observation that light exposure suppresses pineal BOLD activation in a luminance-dependent manner is consistent with the known suppression of melatonin production in response to light (Lewy et al., 1980). This light-induced melatonin suppression is retained following severe degeneration of rods/cones, in both humans (Czeisler et al., 1995) and mice (Lucas et al., 1999). Its spectral tuning is strikingly similar to that of melanopsin, rather than to that of rods or cones (Brainard et al., 2001), and is a common assay for the integrity of the non-image-forming visual system. The similar form of the pineal and prefrontal light-evoked responses, and its divergence from light responses in the visual cortex and LGN, is yet another plausible indicator for ipRGCs contribution to the observed suppression of prefrontal activation by light.

### Other sources of time-locked increases in brain activation

Eye movements or blinking, possibly coinciding with the light stimulus onset, might have induced the observed time-locked activation increase is several brain regions. The cortical areas involved in eye movements control have been reported to include the frontal eye field, supplementary eye field, posterior parietal lobe, dorsolateral PFC which lies in the middle frontal gyrus, and the anterior and posterior cingulate cortex (Pierrot-Deseilligny et al., 2004). Consequently, one of the luxotonic transiently-modulated regions, the middle frontal gyrus, might be associated with eye movements (Fig. 7, T3). Eye blinking might potentially represent another confounding factor.

However, eye blinks have been shown to be associated with activation increases in the anterior cingulate gyrus, right inferior frontal gyrus, right superior temporal gyrus, left superior temporal gyrus, and cuneus (Hanakawa et al., 2008), but not with activation modulation in any of the 17 luxotonic regions identified in the current study as exhibiting a significant transient component (Fig. 7). Taken together, aside from the transient luxotonic regions identified in the middle frontal gyrus, all other regions identified in the current study most likely represent a true luxotonic behavior.

### An overlap between luxotonic regions in which activation showed significant sustained and transient components

While this study focused on the light exposure effect on steady-state brain activation, our analysis identified 17 brain regions exhibiting monotonic responses to luminance levels with a significant transient component (Fig. 7; see Supplementary Text for an overview of the functions mediated by those regions). Our identification of luxonic regions in the above-mentioned cortical and subcortical areas raises the possibility that light might affect at least some of these brain centers, and possibly modulate an array of sensory, motor, emotional, cognitive, and homeostatic functions. Further study into the effect of light exposure on those brain regions and the functions they mediate would shed light on these questions. We have found that sustained and transient responsive regions were almost completely segregated, except for one nearly contiguous strip running along the parieto-occipital sulcus and the dorsal cuneus (Fig. 8). This cuneus region might be in a position to modify information transferred from the primary visual cortex to extrastriate cortices (Vanni et al., 2001).

## Acknowledgments

We would like to thank Rebeca Waugh, a technician at the Brown University neuroimaging facility, and Eli Shmueli for valuable comments on the manuscript. This project was supported by a grant of the National Institute of Psychobiology of Israel, the Banting Postdoctoral Fellowship of Canada to S.S.; NIH grant (R01 EY12793), an Alcon Research Institute Award to D.M.B.; NIH grant (P20GM103645) and the Division of Biology and Medicine, Brown University to J.N.S.

## Author contributions

Conceptualization: SS, DMB, JNS

Methodology: SS, MSW

Investigation: SS, MSW

Visualization: SS, MSW, DDL

Supervision: SS, DMB

Writing—original draft: SS

Writing—review & editing: SS, DMB, JNS, MSW, DDL

## Competing interests

The authors declare no competing interests.

## Data and materials availability

All data are available in the main text or the supplementary materials.

## Lead contact

Further information and requests for resources should be directed to and will be fulfilled by the Lead Contact, Shai Sabbah.

## Materials availability

This study did not generate new unique reagents.

## Data and code availability

Original data for all figures and datasets in this paper, as well as new code generated for this paper are available upon request to the lead contact.

## EXPERIMENTAL MODEL AND SUBJECT DETAILS

We recruited 20 healthy female and male participants from the local community (age 20-30 years old; 24.3 ± 3.7 years, mean ± s.d.). All participants provided written informed consent according to established and approved Institution Review Board guidelines for human participation in experimental procedures at Brown University. We adhered to the principles of the Declaration of Helsinki. Participants received modest monetary compensation.

## METHOD DETAILS

### Functional magnetic resonance imaging (fMRI)

We used BOLD functional MRI (3T Siemens PRISMA, 68 oblique axial slices tilted 30 degrees from the plane of the AC-PC line with 2 mm isotropic voxels, TR 2 sec, TE 30 ms, FOV 212 mm) to explore whole-brain activation patterns of 20 healthy participants in response to full-field diffused light stimuli. Participants underwent five runs, each consisting of three groups, and each group consisting of three blocks. A block contained all four luminance levels in pseudorandom order (10.2, 12.1, 13.1, and 13.8 log photons cm^-2^ s^-1^; 30 sec epochs). Visual stimuli were presented using a digital light processing (DLP) projector (Optoma EP719) coupled with a plano-convex lens that was used to focus the light onto the mirror of the head coil. Participants wore Teflon goggles to diffuse the light and remove any ambient patterned visual information. To monitor the level of alertness, participants performed an auditory discrimination task in which they discriminated between two infrequently presented tones (1 and 2 kHz) and indicated the perceived tone by pressing the left button for the 1 kHz tone and the right button for the 2 kHz tone. Luminance level was measured behind the Teflon goggles at the plane of the participants eyes using a fiber-coupled (core diameter 1000µm, 0.5 NA; Ocean Optics) absolute-irradiance-calibrated spectrometer (USB4000-UV-VIS-ES, Ocean Optics).

We did not pharmacologically control pupil size. Thus, the participants’ pupils likely constricted or dilated depending on stimulus luminance. Increased luminance leads to pupil constriction and restricts the amount of light reaching the retina. However, this occurs within the first 1-2 sec. upon light onset, and is dependent on luminance level (Barbur et al., 1992). This change in pupil diameter should theoretically diminish the light-evoked activation in the LGN and visual cortex, while in our setup, increased luminance led to greater activation in the LGN and visual cortex.

Therefore, whatever was the pupil size at the time of recording, it evidently did not restrict observing the relationships between luminance and activity level, albeit without having an indication of whether the observed activity level represents an absolute maximal value. At most, pupil size might have led to an underestimation of the highest potential activation in response to light.

### Spatiotemporal processing of fMRI data

Non-linear warping to MNI space (using AFNI SSwarper) was used for anatomical alignment. The surviving anatomical detail, including subcortical regions, indicates a robust alignment.

Individual data were slice-time and motion corrected, spatially smoothed (4 mm kernel).

### Identification of luxotonic brain regions

We used a combination of criteria to reveal brain regions whose activation varied monotonically with luminance level and exhibited either a significant transient component or a significant sustained component. Each 30 sec. stimulus block was modeled with a pair of regressors, one transient and one sustained (Horiguchi et al., 2009), using a Generalized Least Squares approach with restricted maximum likelihood estimation of the temporal auto-correlation structure (using the AFNI program (Cox, 1996) with 3dREMLfit). The transient regressor convolved the hemodynamic response function (HRF) with a 1 sec. stimulus event occurring at the transition time from one stimulus intensity to the next, allowing us to capture transient onset and offset responses, known to occur for luminance level increments. Transient responses may be produced both by luminance onsets and by luminance offsets. Because the initial response at each luminance transition includes both the onset response for the current luminance level and an “offset” response for the previous level, an additional parametric regressor was used for the transients. The parametric regressor varied linearly in amplitude according to the absolute value of each luminance increment or decrement; that is, both increments and decrements of luminance were assumed to produce a transient effect of similar magnitude and polarity that varied according to step size. The sustained regressor convolved the HRF with the entire 30 sec. stimulus period and allowed us to estimate sustained light-evoked responses independently of any transient onset or offset responses. Modeling our data using these transient and sustained regressors yielded more stable results compared to modeling with TENT functions at each repetition time (RT) because it involves the fitting of only one parameter (beta coefficient) for each block function, as opposed to fitting separate parameters for each of the 15 individual TENT functions. Finally, a single additional regressor was included to account for auditory stimulation and the corresponding behavioral response. We did not analyze the response to auditory stimuli separately but instead treated it as nuisance effect. A linear contrast was used to identify voxels having monotonically increasing or monotonically decreasing responses according to luminance levels. Group level contrasts were calculated using AFNI’s 3dMEMA program for calculation of mixed effects, taking into account both within and across subject variability with effects calculated based on the coefficients from the single subject contrasts. Clusters were identified using a voxel wise significance of p<0.001 and a family wise error (FWE) of 0.05 (minimum cluster size = 27 voxels). A small volume correction was performed to find the same FWE only for thalamic regions (minimum cluster size = 9 voxels), which did not otherwise survive the full brain false discovery rate (FDR) correction.

Luminance-Response (LR) curves were calculated using the early phase of response (*R*_*0*_) for brain regions that exhibited a significant transient component, and using the late, steady-state, phase of response (*R*_*s*_) for brain regions that exhibited a significant sustained component. *R*_*0*_ and *R*_*s*_ were taken as the mean BOLD activation over the first and last 10 sec. of the 30 sec. visual stimuli, respectively. The sensitivity of a brain region to light was calculated as the difference in *R*_0_or *R*_s_between the second lowest (L2) and highest (L4) stimulus luminance divided by the change in luminance (1.7 log photons cm^-2^ s^-1^). All data were analyzed using custom Matlab scripts.

## Supplemental information

### Supplemental Text

#### A transient effect of light on brain activation

Our analysis identified 17 brain regions in which light affected activation in a luminance-dependent manner but only transiently. These included the precentral gyrus (primary motor cortex, T6, T7), which together with the supplementary motor area (T5) (Penfield and Welch, 1951), paracentral lobule (T11) (Lim et al., 1994) and other structures are involved in the planning, regulation, and execution of voluntary movements. Indeed, light has been shown to play a significant role in shaping neuronal activity in the motor cortex during locomotion (Armer et al., 2013). The frontal operculum (T8, T9) is a critical node in a network controlling activation in other brain areas to perform a wide array of cognitive tasks (Dosenbach et al., 2006; Eichele et al., 2008; Fair et al., 2007). The inferior occipital gyrus (T15) contributes to processing of facial features (Arcurio et al., 2012; Liu et al., 2010). The middle occipital gyrus (T14) processes visual spatial information (Renier et al., 2010). The parietal operculum (T11) is the site of the secondary somatosensory cortex (SII), and thus contributes to perception of vibration, light touch, pain and visceral sensations (Eickhoff et al., 2006). The insular cortex (T12) plays a central role in the regulation of homeostasis, is the site of the primary gustatory cortex, is involved in the perception of visceral sensation, and receives auditory, olfactory, vestibular, and somatosensory input from diverse association areas, as well as the anterior orbitofrontal cortex, PFC, and other limbic structures (Nieuwenhuys, 2012), suggesting its heavy involvement in cognitive and emotional processes.

## Supplemental figures

**Figure S1.**
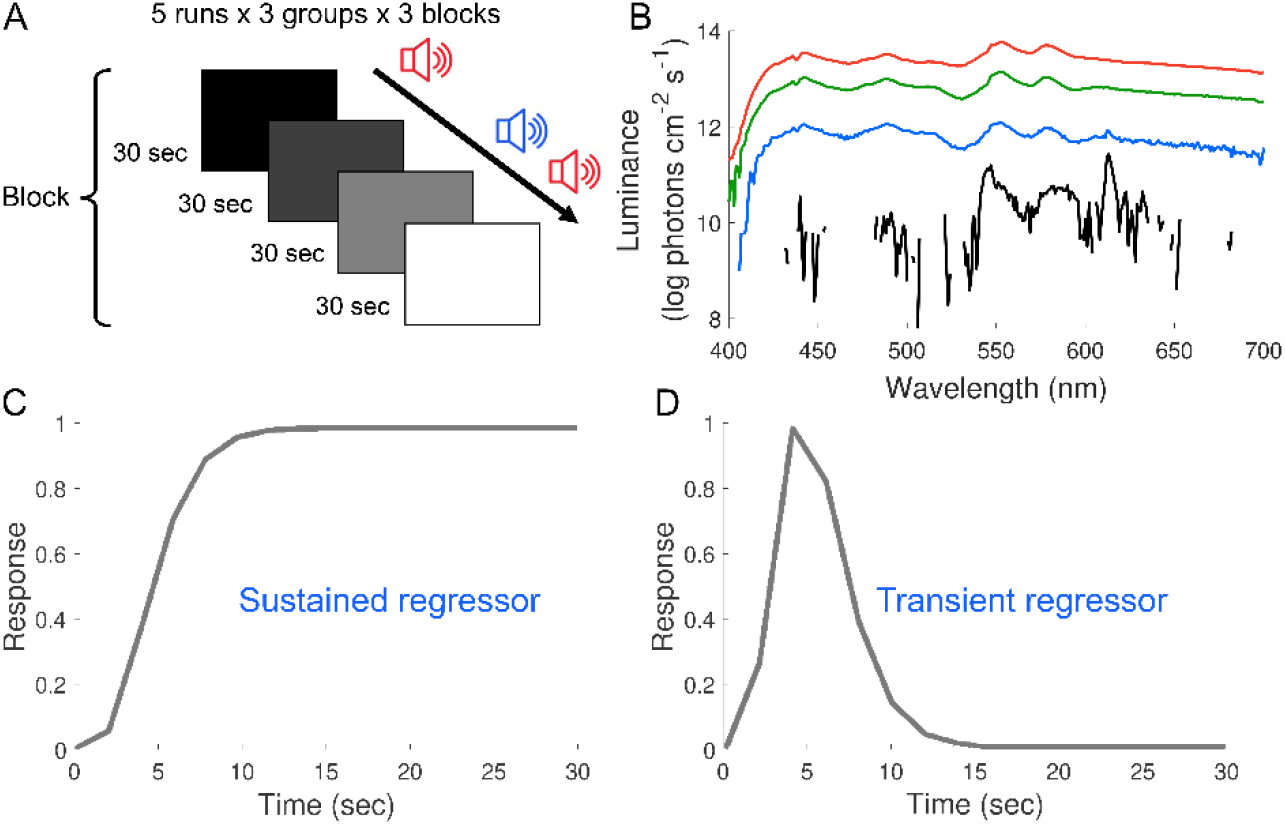
Experimental protocol, stimuli, and analysis. **(A)** Experimental protocol. Each participant underwent five runs, each consisting of three groups, and each group consisting of three blocks (thus, 45 blocks per subject). A block consisted of 30-sec full-field diffused light stimuli at four luminance level levels presented in pseudorandom order. To monitor alertness, participants were asked throughout the protocol to perform an auditory discrimination task of discriminating between two tones, presented in pseudorandom order, by pressing on one of two buttons. **(B)** Spectra of the four visual stimuli. Missing data in the spectra of the lowest luminance level (black) represent spectral ranges in which luminance was too low to be detected by the spectrophotometer. (**C, D**) Each block of four 30 sec. stimuli was modeled with a pair of regressors: (1) a sustained regressor with a 30 sec. stimulus event, covering the entire 30 sec. stimulus period, and (2) a transient regressor with a 1 sec. stimulus event occurring at the transition time from one stimulus luminance to the next.

**Figure S2.**
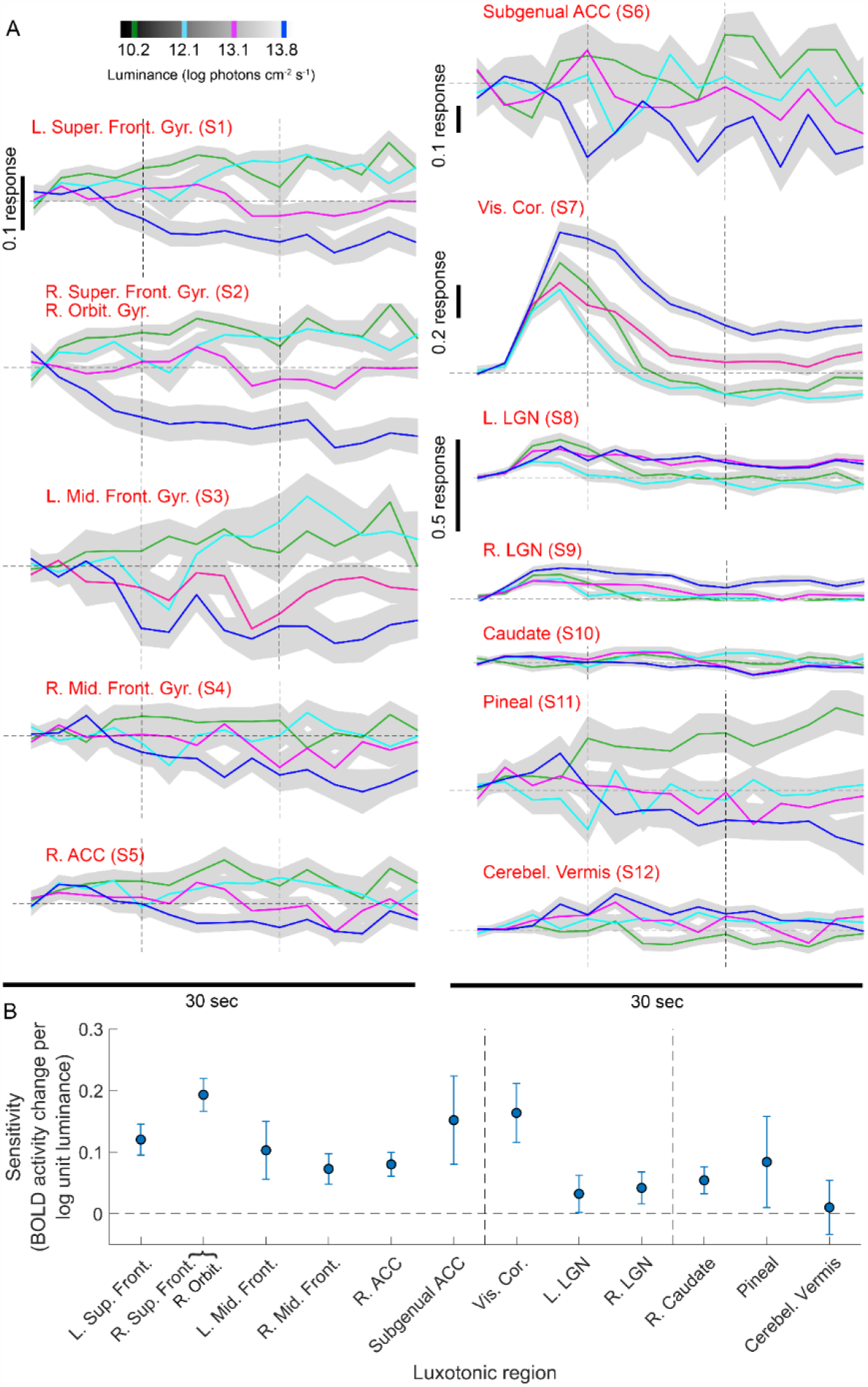
Light-evoked BOLD responses over time and light sensitivity in sustained luxotonic regions. **(A)** BOLD responses over the 30-sec light stimulus, for four luminance levels, for each of the luxotonic regions that exhibited a statistically significant sustained component. Vertical scale bars are related to the panel in which they appear and to all consecutive panels. **(B)** Light sensitivity in cortical regions was higher than in subcortical regions.

**Figure S3.**
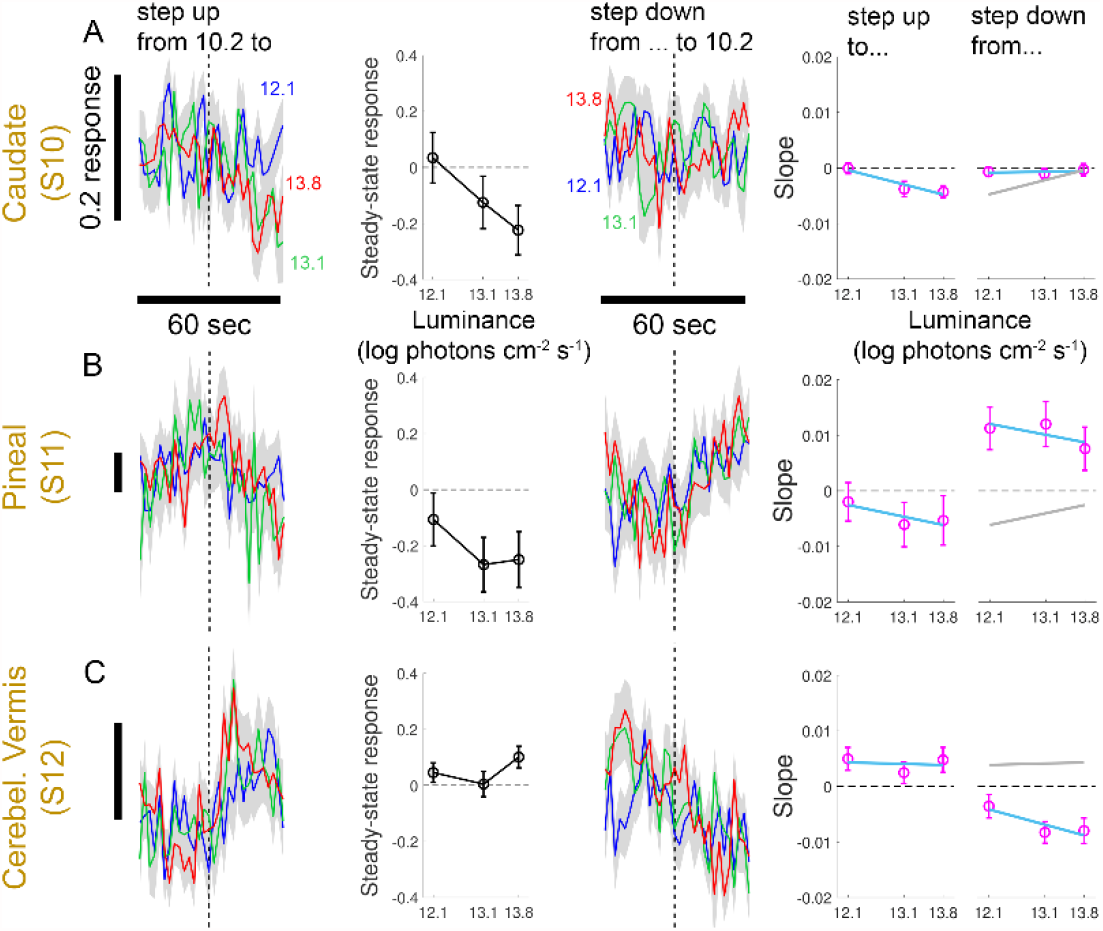
Light monotonically varies the activity in the pineal, caudate and cerebellar vermis. Conventions as in Figs. 3 and 4.

**Figure S4.**
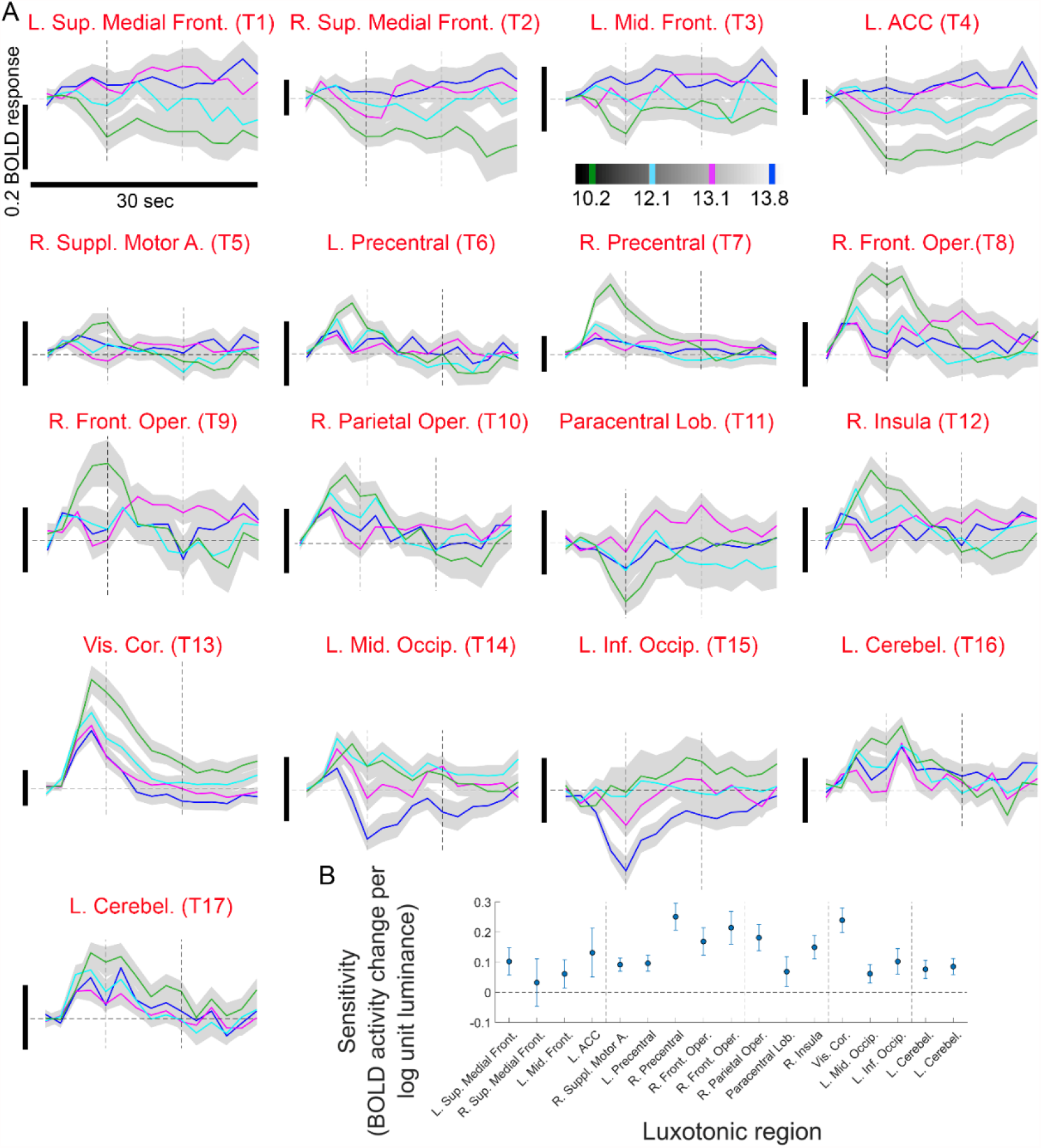
Light-evoked BOLD responses over time and light sensitivity in luxotonic regions with a transient component. **(A)** BOLD responses over the 30-sec light stimulus, for four luminance levels, for each of the 17 luxotonic regions that exhibited a statistically significant transient component. Vertical scale bars represent 0.2 BOLD response. **(B)** Light sensitivity in regions of the visual cortex and operculum were highest.

**Figure S5.**
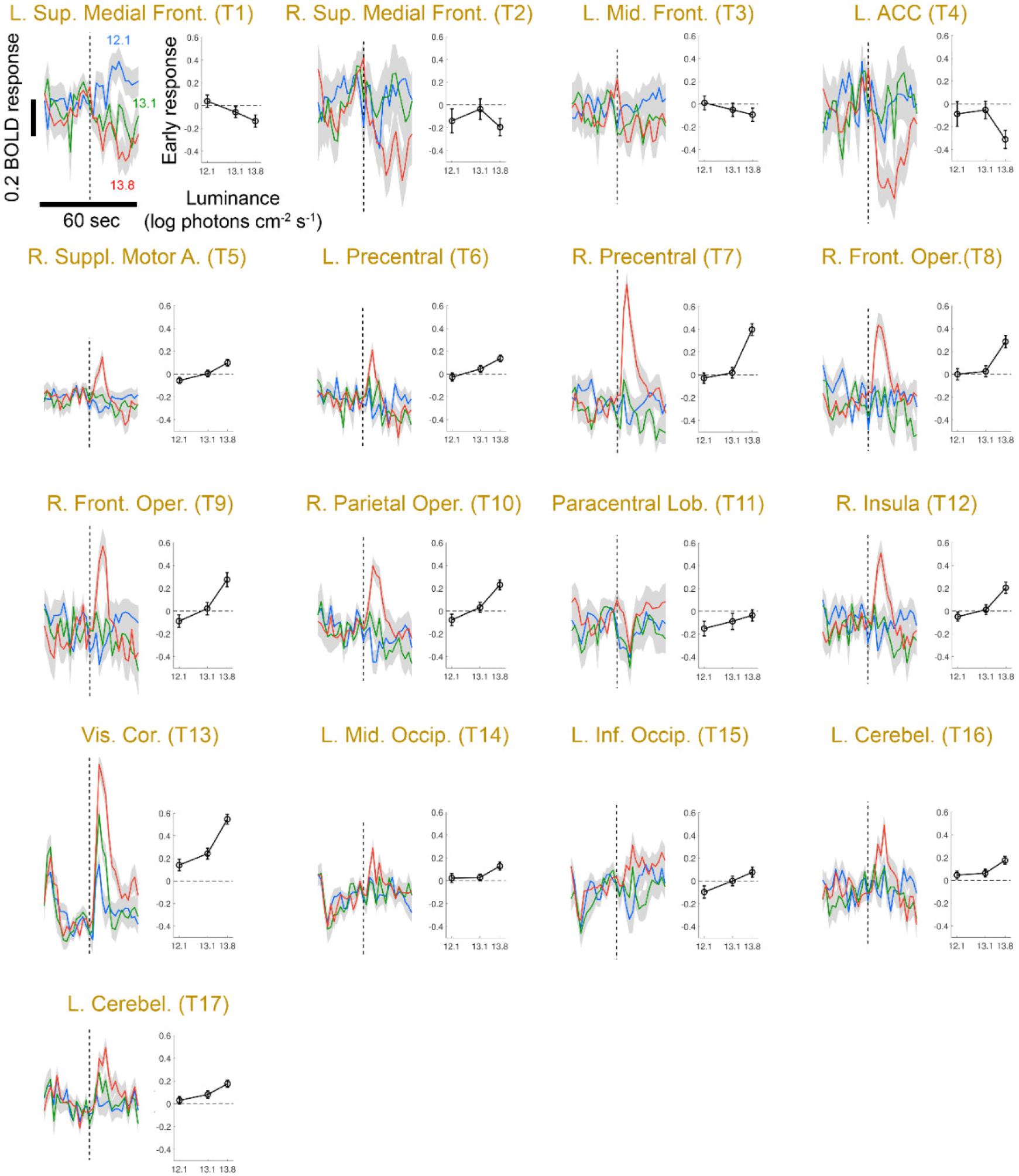
Light transiently modulates early cortical and cerebellar activity. (Left) BOLD Light-evoked responses to step-up transitions – transitions in which luminance was stepped up from the same, lowest, luminance level (L1, 10.2 log photons cm^-2^ s^-1^). Responses around individual transitions are depicted in different colors. Shaded gray areas represent the standard error from the mean. Dashed vertical line marks the transition time. (Right) Early prefrontal activity monotonically decreased with increasing luminance level (regions T1-T4) while early activity in all other regions (T5-T17) increased with luminance level. Vertical bar, representing 0.2 BOLD response, applies to all panels.

## Notes

### Competing Interest Statement

The authors have declared no competing interest.

